# Extensive Phenotypic Changes Associated with Large-scale Horizontal Gene Transfer

**DOI:** 10.1101/001362

**Authors:** Kevin Dougherty, Brian A. Smith, Autumn F. Moore, Shannon Maitland, Chris Fanger, Rachel Murillo, David A. Baltrus

## Abstract

Horizontal gene transfer often leads to phenotypic changes within recipient organisms independent of any immediate evolutionary benefits. While secondary phenotypic effects of horizontal transfer (i.e. changes in growth rates) have been demonstrated and studied across a variety of systems using relatively small plasmid and phage, little is known about how size of the acquired region affects the magnitude or number of such costs. Here we describe an amazing breadth of phenotypic changes which occur after a large-scale horizontal transfer event (~1Mb megaplasmid) within *Pseudomonas stutzeri* including sensitization to various stresses as well as changes in bacterial behavior. These results highlight the power of horizontal transfer to shift pleiotropic relationships and cellular networks within bacterial genomes. They also provide an important context for how secondary effects of transfer can bias evolutionary trajectories and interactions between species. Lastly, these results and system provide a foundation to investigate evolutionary consequences in real time as newly acquired regions are ameliorated and integrated into new genomic contexts.

## INTRODUCTION

<u>H</u>orizontal <u>G</u>ene <u>T</u>ransfer (HGT), the movement of genetic material between individuals without reproduction, is a major evolutionary force within microbial communities and impacts genome dynamics across all all life (Keeling, 2009; Syvanen, 2012). Although HGT events often provide direct fitness benefits to recipient cells, such as antibiotic resistance, integration of foreign DNA is an inefficient process (Dahlberg and Chao, 2003; Diaz-Ricci and Hernández, 2000; Park and Zhang, 2012). As a result, newly acquired regions often interfere with physiological, genetic, and regulatory pathways to cause changes independent of phenotypes under immediate or direct selection pressures (Baltrus, 2013). Numerous studies have demonstrated the existence of such costs by documenting changes to fitness, growth rate, or other phenotypes after the transfer of relatively small genomic regions. However, little is known about how the potential for and diversity of HGT-associated costs scales with size of the transferred region.

A variety of non-mutually exclusive mechanisms potentially contribute to costs of HGT. For instance, recently acquired genes are typically expressed at inefficient levels leading to limitations in resources such as ribonucleotides, amino acids, or ATP (BRAGG and WAGNER, 2009; Stoebel et al., 2008). Additional genes can occupy molecular machines required for basic cellular functions, such as polymerases and ribosomes, and sequester these limiting enzymatic resources from more critical activities (Dethlefsen and Schmidt, 2007; Shachrai et al., 2010). Foreign proteins may not fold correctly in their new cellular contexts, which could lead to disruption or triggering of stress responses (Drummond and Wilke, 2008; Park and Zhang, 2012). Recently acquired regions may disrupt flux through cellular systems, leading to the buildup of toxic intermediates (Gonçalves et al., 2011; Shintani et al., 2009). While such costs have been directly observed in laboratory experiments, retrospective studies across genomes add an additional layer of complexity as there exists an inverse correlation between gene retention after HGT and number of protein-protein interactions affected (Cohen et al., 2011). In most cases the precise molecular mechanisms underlying observed costs of HGT have not been identified, however, both the magnitude and molecular basis for costs could be greatly affected by both the size and gene content of the acquired region.

Costs of HGT have typically been studied by focusing on phenotypic changes after HGT of relatively small plasmids and lysogenic phage, even though large-scale transfers (>60,000 kb) occur at appreciable rates throughout bacteria (Baltrus, 2013; Harrison et al., 2010; Platt et al., 2012; Smillie et al., 2010). We have developed an experimental system to investigate the costs of large-scale HGT by taking advantage of a a ~1Mb megaplasmid which is self-transmissible throughout *Pseudomonas* species (Romanchuk et al, *in prep*). Transfer of this megaplasmid occurs in both liquid and solid media and requires a type IV secretion system. This HGT event introduces approximately 700 ORFs into recipient cells, including many “housekeeping” genes as well as almost a full complement of tRNA loci as is characteristic of megaplasmids (Baltrus et al., 2011; Harrison et al., 2010). Importantly, it does not appear as though full pathways are present for megaplasmid encoded housekeeping gene pathways, so function very likely requires direct interaction with chromosomal networks.

In a parallel manuscript (Romanchuk et al., *in prep*), we demonstrate that acquisition of this megaplasmid lowers fitness of *Pseudomonas stutzeri* by ∼20% and here we report on multiple additional phenotypes affected by large-scale HGT. Specifically, we find that megaplasmid acquisition leads to sensitivity to quinolone antibiotics, DNA intercalating agents, temperature, and killing by other bacterial species. We further find that HGT changes bacterial behavior in that biofilm formation is decreased and motility is increased. This widespread pleiotropy is unprecedented, both in number and diversity of phenotypic changes, and could signal that multiple phenotypic costs occur throughout transfer events (Baltrus, 2013). Moreover, that such pleiotropic relationships between phenotypes are mediated at a systems level by single HGT events creates a unique situation where phenotypic evolution occurs as a by-product of evolutionary amelioration after transfer rather than direct selection on phenotypes themselves. In sum, we document the significant potential for secondary effects of HGT to alter phenotypic evolution and adaptive trajectories across microbial populations. This system further underscores the indirect power of costs of HGT to rapidly generate phenotypic diversity across closely related bacteria.

## RESULTS

### Megaplasmid Acquisition Decreases Thermal Tolerance

Our initial observations suggested that, although we were successfully able to select for conjugation of pMPPla107 from *P. syringae* pv. *lachrymans* 107 to *P. stutzeri* DBL332 when plated out on selective antibiotics at 27°c, conjugations failed when selected at 37°c (data not shown). We have since demonstrated, as one can see in Figure 1A, megaplasmid acquisition sensitizes *P. stutzeri* to growth at 37°C and greater. Although both *P. stutzeri* DBL332 and DBL390 grow well at either temperature, independently created strains containing pMPPla107 (DBL365 and DAB412) appear stressed at 27°C and fail to grow at 37°c (Figure 1A). Similar results were observed for this assay with an additional *P. stutzeri* strain (DBL408) which acquired the megaplasmid independently of DBL365 (data not shown). To further quantify this effect, we measured how changes in temperature alter competitive fitness for two related megaplasmid containing strains (DBL365 and DBL453). As one can see in figure 1B, presence of pMPPla107 decreases competitive fitness by 22% and 12% for DBL365 and DBL453 respectively at 35°c compared to 27°c. Analyzed within a full factorial ANOVA framework, this effect of temperature is significant (F_1,3_=45.33, p = 0.0067). Furthermore, analyzed as contrasts within the ANOVA framework, each strain’s fitness is significantly lower at 35°C compared to 27°C (p < 0.05). Since DBL453 is derived from DBL365 by recombining out the tetracyline marker, these results are not due to the marker itself. Therefore, two separate assays confirm that large-scale HGT of a megaplasmid decreases thermal tolerance in *P. stutzeri*.

**Figure 1.**
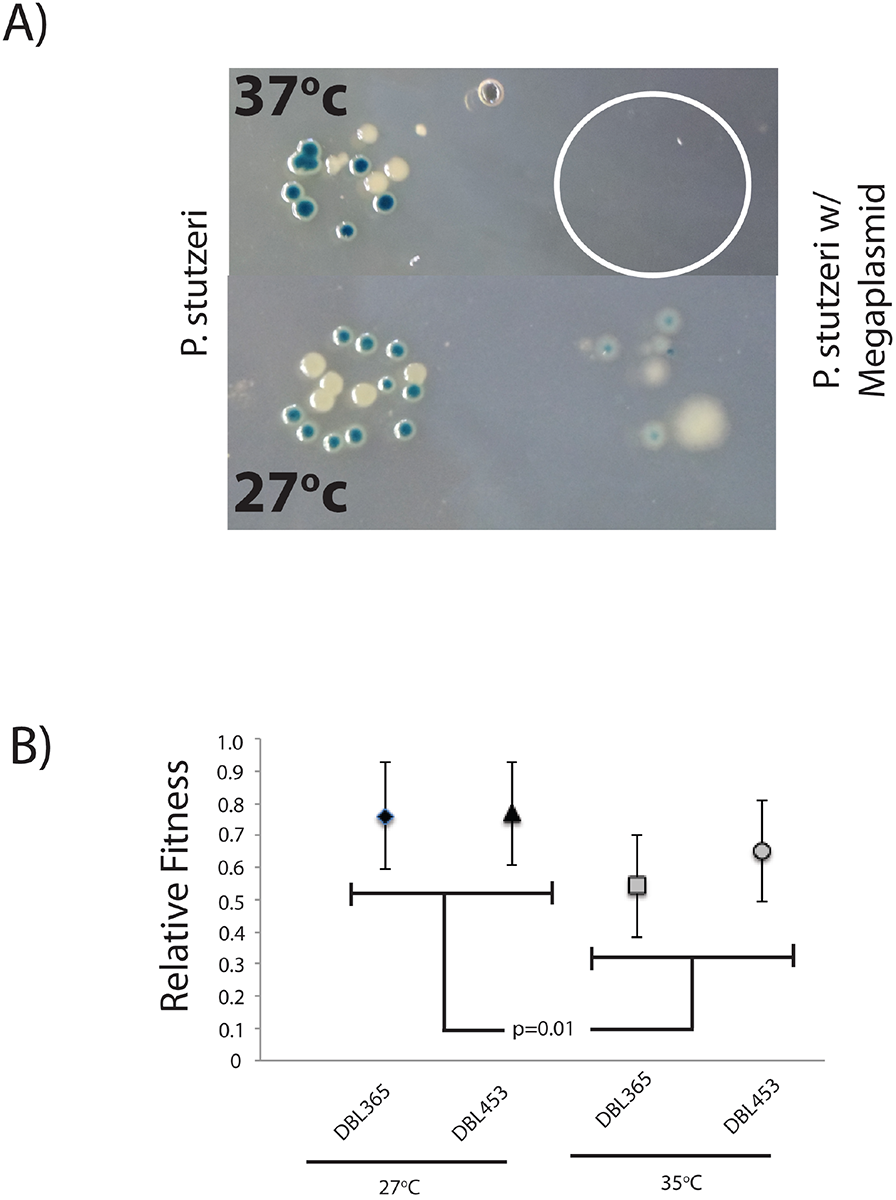
Megaplasmid Acquisition Decreases Thermal Tolerance. A) A dilution series of strains DBL332/DBL365 (white) and DBL390/DBL412 (blue), which lack/ contain megaplasmid pMPPla107 were plated at either 27°C or 37°C. *P. stutzeri* normally grows at both temperatures (left), but strains containing the megaplasmid (right) only grow at 27°C. Pictures were taken at the lowest dilution with growth (approximately 1:1 × 10^9^). White circle reflects where megaplasmid containing strains were spotted but didn’t grow. B) Competitive fitness assays demonstrate that fitness of megaplasmid containing strains is significantly lower at 35°C compared to 27°C (p = 0.01). Fitness is normalized so that *P. stutzeri* lacking the megaplasmid is 1, error bars shown are ± 2 standard errors.

### Megaplasmid Acquisition Decreases Antibiotic Resistance

We used Biolog assays (Bochner, 2003) to identify phenotypic changes associated with acquisition of megaplasmid pMPPla107 by *P. stutzeri* DBL332 (File S1). Overall, aside from an overall consistent negative effect of pMPPla107 on grown of *P. stutzeri,* acquisition of the megaplasmid only significantly changed a handful of phenotypes across replicates (File S1). One striking result is that megaplasmid acquisition decreases resistance of *P. stutzeri* to a variety of quinolone antibiotics as well as DNA intercalating agents such as 7-hydroxycoumarin. We followed up these results by performing replicated growth curves in both Naldixic acid and 7-hydroxycoumarin over a series of different concentrations. As shown in Figures 2A and B, megaplasmid presence lowers the minimal inhibitory concentration of *P. stutzeri* to both 7-hydroxycoumarin and Naldixic acid, thus replicating the results from the Biolog assay. Similar results were observed for this assay with an additional *P. stutzeri* strain (DBL408) which acquired the megaplasmid independently of DBL365 (data not shown). To further quantify this difference in antibiotic resistance, we measured competitive fitness between DBL332 and DBL365 in 0 and 4μg/mL Naldixic acid. As one can see in figure 2C, presence of pMPPla107 decreases competitive fitness by 31% and 28.5% for DBL365 and DBL453 respectively in the presence of 4μg/mL Naldixic acid. Analyzed within a full factorial ANOVA framework, the effect of antibiotic is significant (F_1,2_=20.7564, p = 0.045). Furthermore, analyzed as contrasts within the ANOVA framework, each strain’s fitness is significantly lower at 4μg/mL Naldixic acid compared to standard SW-LB (p < 0.05). Since DBL453 is derived from DBL365 by recombining out the tetracyline marker, these results are not due to the marker itself Therefore, two separate assays confirm that large-scale HGT of a megaplasmid decreases resistance to quinolone antibiotics in *P. stutzeri*.

**Figure 2.**
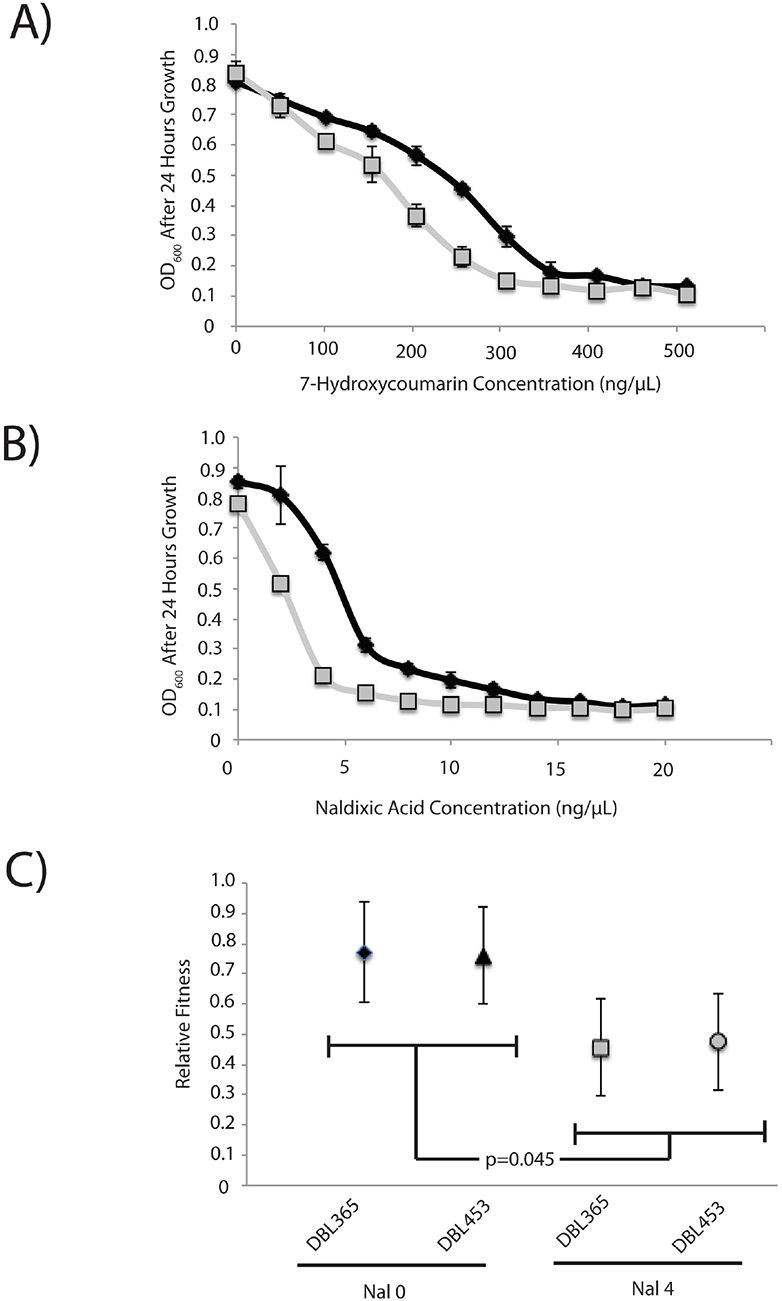
Megaplasmid Acquisition Decreases Resistance to 7-hydroxycoumarin and Naldixic acid. A) A dose response curve comparing bacterial growth (OD_600_) of bacterial strains which contain (grey, DBL365) or lack (black, DBL332) the megaplasmid pMPPla107 across 7-hydroxycoumarin concentration. Curve is representative of at least 3 different assays. B) A dose response curve comparing bacterial growth (OD_600_) of bacterial strains which contain (grey, DBL365) or lack (black, DBL332) the megaplasmid pMPPla107 across Naldixic acid concentration. Curve is representative of at least 3 different assays. C) Competitive fitness assays demonstrate that fitness of megaplasmid containing strains is significantly lower in 4 µg/mL Naldixic acid compared to 0 µg/mL (p = 0.045). Fitness is normalized so that *P. stutzeri* lacking the megaplasmid is 1, error bars show ± 2 standard errors in all cases.

### Megaplasmid Acquisition Decreases Biofilm Formation

We witnessed that, after extended periods of growth in liquid media, *P. stutzeri* DBL332 forms a mass at the bottom of pipette tip within the culture after approximately 2-4 days (Figure 3A and B). This mass remains regardless of how long cultures are left to incubate (longest tested period was 10 days, data not shown). However, strain DBL365 does not form such a mass, regardless of how long the culture is incubated for (data not shown). To quantify this effect, we grew replicate cultures of strains DBL332 and DBL365 in SWLB media containing a single pipette tip. As one can see in figure 3C, even though population sizes of bacterial population suspended in liquid media are roughly equivalent (6.6 × 10^9^ and 4.8 × 10^9^ CFU/mL for DBL332 and DBL365 respectively), population sizes of bacteria attached to the pipette tip are much greater in the strain that lacks the megaplasmid (1.4 × 10^8^ compared to 7.6 × 10^6^). Therefore, whereas roughly 2% of *P. stutzeri* forms a mass on the pipette tip in the absence of pMPPla107, only 0.15% of the population forms this mass in a strain containing the megaplasmid. Similar results were observed for this assay with DBL453 as well as an additional *P. stutzeri* strain (DBL408) which acquired the megaplasmid independently of DBL365 (data not shown). We also note that we attempted to perform traditional biofilm assays (O’toole et al., 2011) to compare strains that contain or lack the megaplasmid, but slower growth of the megaplasmid strains made comparisons difficult to interpret (data not shown).

**Figure 3.**
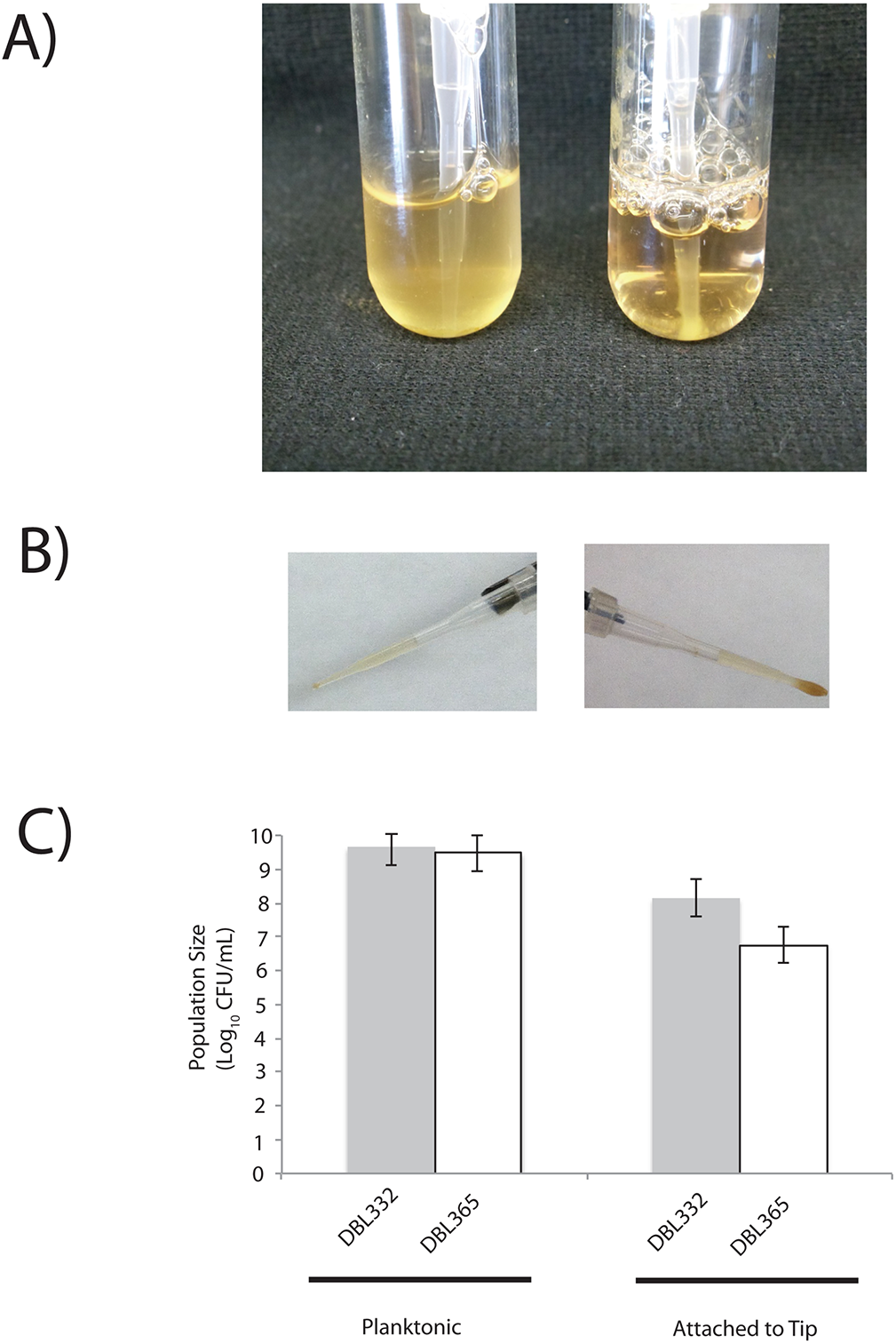
Megaplasmid Acquisition Decreases Biofilm Formation. A) Four day old cultures of strains which lack (DBL332, right) or contain (DBL365, left) megaplasmid pMPPla107 grown in M9 media supplemented with 10Mm succinate. B) Pipette tips harvested after four days of growth in 2mL SWLB media cultures for either DBL365 (left) and DBL332 (right). C) Bacterial population sizes of DBL332 (grey) and DBL365 (white) after four days of growth in 2mL cultures of SWLB media. Population sizes of planktonic bacteria are shown on the left, while those harvested from pipette tips are shown on the right. Error bars show ± 2 standard errors in all cases.

### Megaplasmid Acquisition Increases Motility

We tested whether megaplasmid acquisition altered motility or chemotaxis in *P. stutzeri* using standard assays (Hockett et al., 2013). Briefly, 1uL of liquid culture was plated into semisolid agar and strains were placed into the incubator. After two days, size of the bacterial halo surrounding the inoculation point was quantified. As shown in figure 4A, strains containing the megaplasmid consistently had larger halos than strains that lack the megaplasmid. Quantification shows that this increase in halo size is approximately 27% (F_1,4_=59.294, p = 0.0015). Similar results were observed for this assay with DBL453 as well as an additional *P. stutzeri* strain (DBL408) which acquired the megaplasmid independently of DBL365 (data not shown).

**Figure 4.**
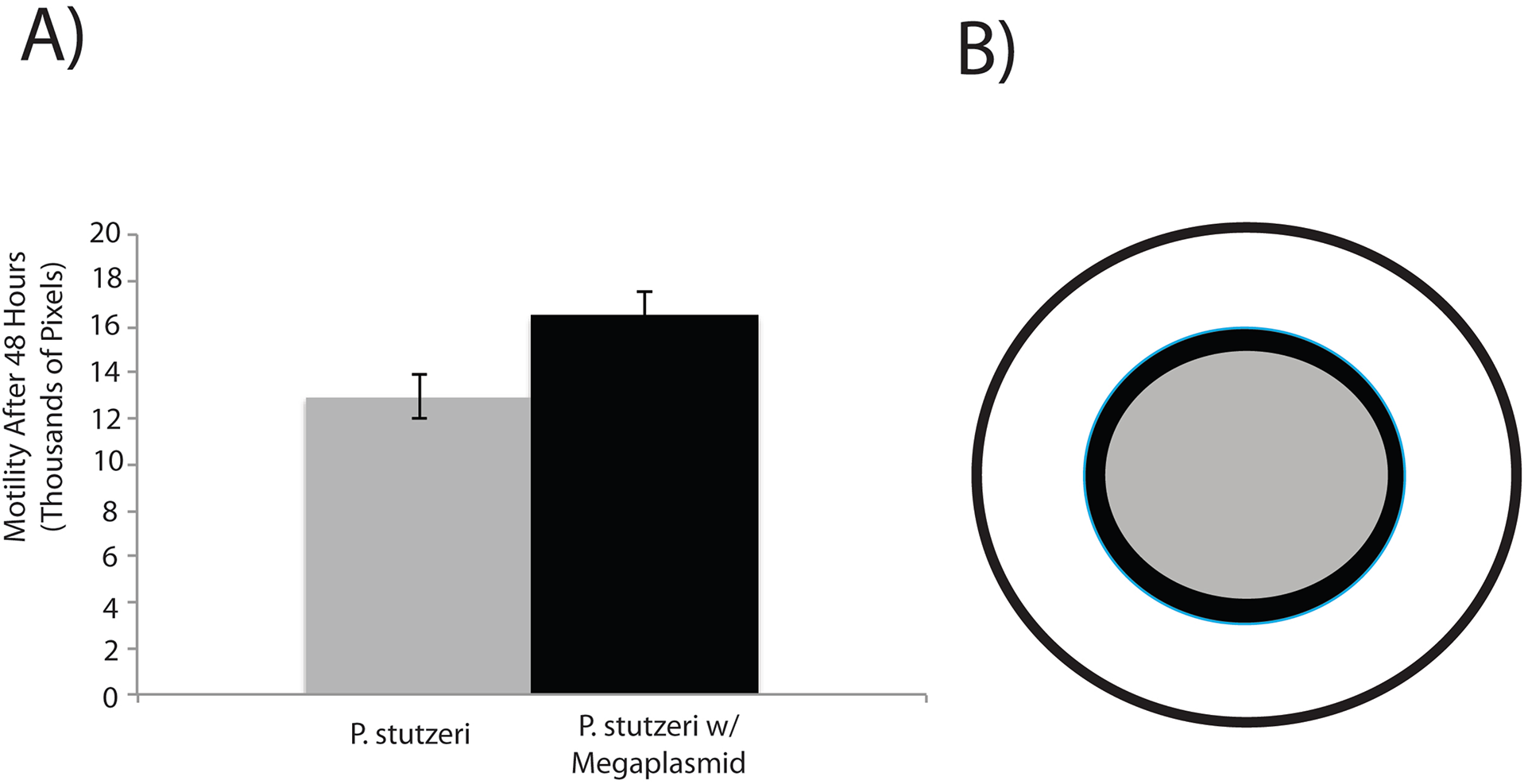
Megaplasmid Acquisition Increases Motility or Chemotaxis. A) Halo size in soft agar is significantly larger for bacterial strains which contain the megaplasmid (DBL365, black) compared to those that lack it (DBL332, grey). Error bars show ± 2 standard errors. B) Average difference in halo size represented in circular form. Grey circle is *P. stutzeri* while black circle represents *P. stutzeri* containing the megaplasmid.

### Megaplasmid Acquisition Increases Sensitivity to Supernatants from Other Bacterial Species

*P. aeruginosa* strains are known to produce a variety of antimicrobial products during growth within laboratory culture, including pyocins and quinolones (Heeb et al., 2010; Inglis et al., 2012; Singh et al., 2010). Since interactions between bacterial species occur frequently within the environment, and are essential for HGT of the megaplasmid to occur, we tested for whether megaplasmid acquisition altered sensitivity of *P. stutzeri* to *P. aeruginosa* supernatant. We found that supernatant from *P. aeruginosa* interfered with growth of *P. stutzeri* cultures, but only when strains contained the megaplasmid (Figure 5). Similar results were observed for this assay with DBL453 as well as two additional *P. stutzeri* strains (DBL408) and *P. syringae* pv. *lachrymans* strain (DAB837) which both acquired the megaplasmid independently of DBL365 (data not shown). Therefore, megaplasmid acquisition alters interactions between bacterial species by sensitizing strains to killing by other bacterial species during normal growth.

**Figure 5.**
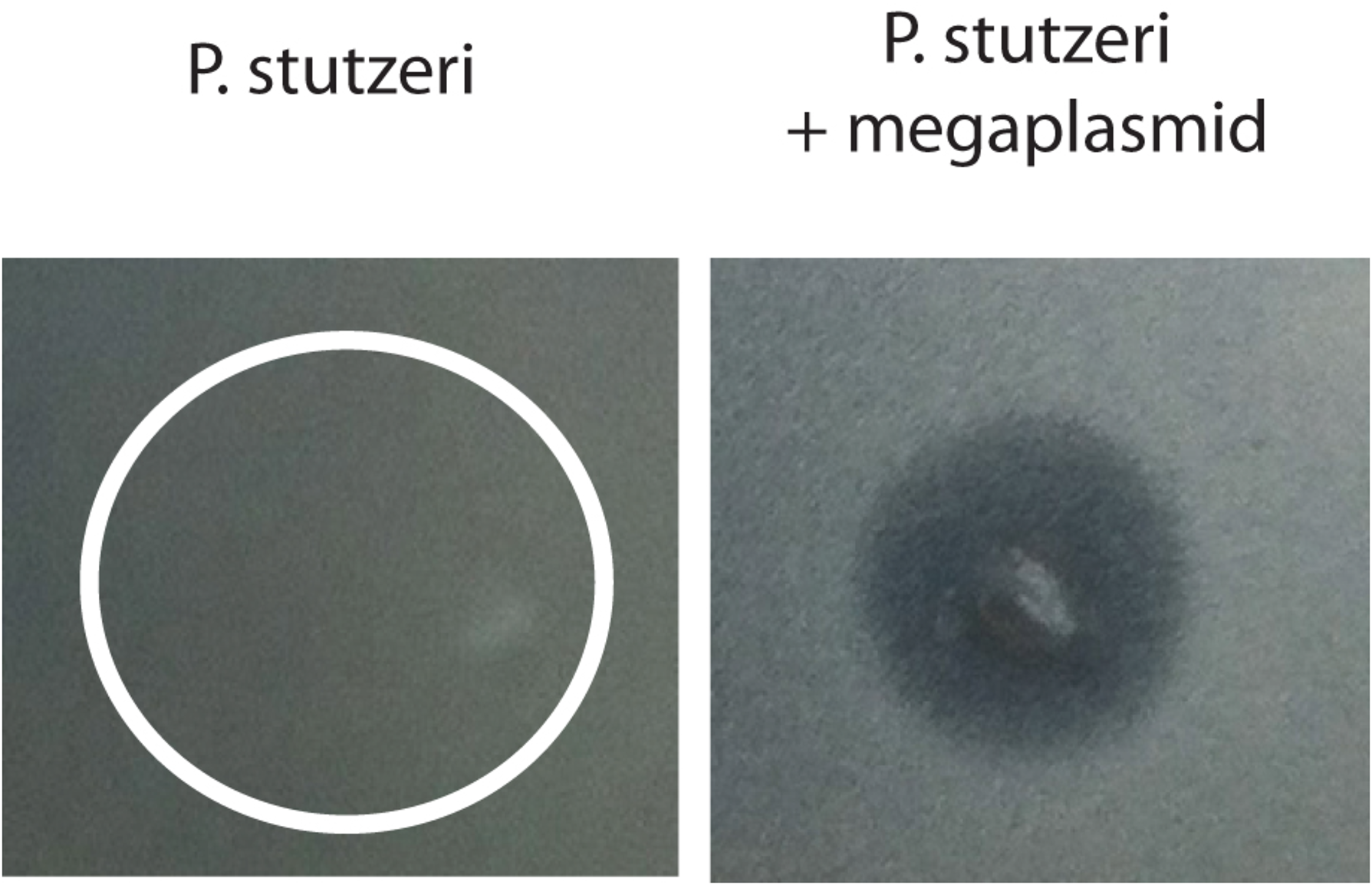
Megaplasmid Acquisition Increases Sensitivity to *P. aeruginosa* Supernatant. Purified supernatant from *P. aeruginosa* PA14 was spotted on lawns of *P. stutzeri* that either lacked (DBL332, left) or contained (DBL365, right) megaplasmid pMPPla107. White circle on left panel represents where supernatant was spotted onto lawn where *P.stutzeri* continued to grow. Right panel shows clearing area in laws where supernatant was spotted. Photo is representative of >3 independent experiments.

## DISCUSSION

For horizontally transferred regions to be maintained over time, they must provide a large enough benefit to avoid loss due to selection or genetic drift. That this benefit may be the primary target of strong selective pressures within a given environment, as with antibiotic resistance, doesn’t preclude the existence of neutral secondary phenotypic changes or HGT-associated costs which are deleterious in other environments (Baltrus, 2013). Although such costs of HGT appear to be widespread, there have been few efforts to investigate how these costs scale with size of the region transferred. Furthermore, even when costs are observed, measurements are often limited to single phenotypes even though multiple cellular systems could be affected (Baltrus, 2013; Dahlberg and Chao, 2003; Gaillard et al., 2008; Heuer et al.; Sato and Kuramitsu, 1998; Shintani et al., 2009). Here we explore a system where HGT increases bacterial genome size by ∼20%, using a megaplasmid which is self-transmissible throughout pseudomonads (Baltrus et al., 2011) and Romanchuk et al., *in prep*). We report that megaplasmid acquisition alters numerous phenotypes within *P. stutzeri* in unprecedented ways, which highlights the potential for large-scale transfers to shape evolutionary dynamics within natural populations. This system provides a unique foundation to explore how evolution affects pleiotropic interactions between phenotypes altered as a result of large-scale HGT events, but also a powerful model to dissect individual interactions underlying these costs at a molecular level.

In a parallel manuscript (Romanchuk et al., *in prep*), we have demonstrated that megaplasmid acquisition by *P. stutzeri* impacts competitive fitness and bacterial growth under standard laboratory conditions. Here we show that megaplasmid acquisition is also accompanied by secondary changes to a variety of phenotypes affecting cellular physiology, environmental survival, and interactions with other species. For instance, megaplasmid acquisition increases sensitivity to quinolone antibiotics as well as stresses such as heat. That these responses are specifically affected by the test environments, as opposed to correlated effects on slower growth across a range of conditions, is highlighted by lack of sensitivity to a variety of other tested conditions (File S1). Surprisingly, we also demonstrate that megaplasmid acquisition also increases sensitivity to a substance present within the supernatant of *P. aeruginosa* cultures. Although we currently do not know the molecule(s) responsible for this effect, multiple bacteriocins and other antimicrobial targets known to be found within the supernatant will be the target of future studies (Heeb et al., 2010; Inglis et al., 2012). Moreover, *P. aeruginosa* is the only pseudomonad that we have failed to be able to conjugate this megaplasmid into (data not shown), so that it appears that costs of HGT could potentially limit horizontal transfer across species. We have further shown that acquisition of the megaplasmid increases bacterial motility (or lowers the chemotaxis threshold) within soft agar and decreases biofilm formation in liquid culture. It is important to note that all of these phenotypic changes are effectively neutral under the defined laboratory conditions which we use to select for successful conjugation because this megaplasmid is engineered to provide tetracycline resistance. Therefore, under selective conditions in the lab, cells that can’t acquire the megaplasmid will die due to antibiotic selection. While selective pressures in nature are likely more complex, these secondary changes could have dramatic effects on ecological strategies or niches between closely related bacterial populations.

This megaplasmid is representative of other large-scale gene transfers in terms of coding capacity, genetic content, and divergence from the recipient genome (Baltrus et al., 2011; Harrison et al., 2010; Smillie et al., 2010). In spite of data demonstrating a negative bias for retention of highly connected genes after HGT events, megaplasmids often contain and can transfer numerous housekeeping genes (Harrison et al., 2010). The megaplasmid within our system itself contains 38 tRNA loci, polymerase subunits, DNA recombination and repair systems, a putative ribosomal protein, as well as other proteins which could be involved in housekeeping functions (Baltrus et al., 2011). At the moment we don’t know whether HGT associated costs are dependent upon any of these genes interfering with chromosomal pathways. However, results obtained herein could represent general outcomes after HGT events or may only be emergent properties of HGT by specific types of larger vectors like chromids.

One major question arising from these results concerns the independence of phenotypic shifts after HGT. Do all observed changes result from a single protein-protein interaction, numerous individual detrimental interactions, or the disruption of complex and interwoven regulatory networks? Furthermore, is the breadth of altered phenotypes a general property of HGT events as a whole or does the number of changes increase with size or gene content of transferred region? One candidate pathway does stand out as a potential mediator of these phenotypes *a priori*. Acquisition of the megaplasmid brings with it hundreds of new genes, the protein products of many of which are membrane localized (Baltrus et al., 2011). Since cell size is limited, the incorporation of additional membrane bound proteins likely disrupts molecular signatures and dynamics of the membrane and could easily trigger or disrupt the envelope stress response (Raivio and Silhavy, 2001). The envelope stress response is conserved throughout bacteria, and responds to a variety of membrane stresses through the action of proteases, anti-sigma factors, as well as a host of other regulators. Since membrane integrity is critical for bacterial survival, the envelope stress response often sits at the top of regulatory cascades which control numerous phenotypes (Balasubramanian et al., 2012). That previous results suggest quinolone sensitivity and sensitivity to *P. aeruginosa* supernatant are correlated with cell membrane integrity provides support for this model (Campos et al., 2006; Hawkey, 2003; Uratani and Hoshino, 1984). Furthermore, chaperone function links both heat and envelope stress responses, since one of the main determinants for both pathways is improperly folded proteins (Jürgen et al., 2010; Kohanski et al., 2008; Raivio and Silhavy, 2001). Lastly, motility and biofilm formation are directly regulated by the envelope stress response in *Pseudomonas*, as both the flagellum and pili are critical membrane bound structures (Baynham et al., 2005; Tart et al., 2006). Alternatively, a variety of independent physiological and regulatory pathways could be responsible for these changes including nutrient limitation triggering the stringent response or the disruption of multiple quorum sensing pathways altering regulation across the genome (van Delden et al., 2001).

Outside of any direct fitness benefits from HGT, that multiple phenotypic changes are linked through single HGT events could skew evolutionary dynamics in unpredictable ways. For example, it is well known that adaptive trajectories and “evolvability” can be influenced by the order that beneficial or compensatory mutations fix (Tenaillon et al., 2012; Woods et al., 2011). Secondary effects of HGT could bias future evolutionary paths within populations by altering magnitude or direction of epistatic interactions between adaptive mutations (Chou et al., 2011). Furthermore, since recently acquired regions often function sub-optimally, the total number of potential beneficial mutations within an environment could be increased due to compensatory changes for the HGT event (Lind et al., 2010). This influx of beneficial mutations correlated to HGT could impact both the order and magnitude of fixed adaptive mutations through clonal interference (Lee and Marx, 2013). Along these lines, significant differences in evolutionary potential could arise based on, for instance, whether antibiotic resistance is introduced through HGT or *de novo* mutation. Such a balance could be further impacted in positive or negative ways by environment specific feedbacks on costs of HGT (Kishony and Leibler, 2003).

Systems biology has made incredible inroads into understanding how genetic, physiological, and regulatory networks structure traits within microbes. Indeed, we are at the cusp of being able to predict and accurately model the phenotypic responses of well-studied bacteria to specific stimuli (Conrad et al., 2011; Durot et al., 2009). However, the value of these models for evolutionary prediction depends on a thorough understanding of how cellular networks interpret and respond to perturbations over evolutionary time scales, including mutation and HGT (Monk et al., 2013; Oberhardt et al., 2009; Pál et al., 2005). Acquisition of a megaplasmid creates new pleiotropic interactions within *P. stutzeri* and changes phenotypic relationships between traits compared to closely related strains. If multiple phenotypic changes are linked through single HGT events, HGT can create pleiotropic relationships between traits that might not otherwise exist. Such context dependence could place firm limits on both accuracy and precision of models for certain bacterial traits outside of controlled laboratory conditions (Monk et al., 2013).

Looking forward, amelioration of costs of HGT could lead to dramatic phenotypic differences between strains descended from a recent common ancestor due strictly to differences in paths of evolution and pleiotropic relationships from HGT (Cooper and Lenski, 2000). These pleiotropic relationships could be reinforced or disrupted depending on which suites of mutations fix over time. For instance, amelioration of megaplasmid associated costs could produce one cluster of strains that is phenotypically indistinguishable from the non-megaplasmid ancestor while another cluster compensates only for costs at 27°c and is unable to grow at 37°C. Furthermore, as a result of costs of the megaplasmid to *P. stutzeri*, increased resistance to quinolone antibiotics such as naldixic acid or ciprofloxin could evolve solely due to compensation for the megaplasmid in the absence of direct selection by quinolones.

Since HGT introduces foreign genes and pathways into novel genomic contexts, each transfer event brings with it great potential to disrupt existing genetic and physiological networks within the recipient cell. Although numerous results have analyzed and dissected costs after the transfer of relatively small plasmids and phage, we have developed a model system with which to explore phenotypic costs associated with large-scale HGT (∼20%). The phenotypic shifts we see are dramatic and include changes of bacterial tolerance to numerous stresses as well as alterations of behavior. Individually, a subset of these changes have been observed in other systems as a result of HGT or as a pleiotropic result of *de novo* adaptive mutations. However, taken as a whole, these results highlight the power of epistasis between the recipient genome and recently acquired regions to completely shift genetic and phenotypic expectations between closely related organisms. Furthermore, our results demonstrate the amazing breadth of phenotypes potentially affected by one HGT event and emphasize how singular evolutionary events can re-wire, reshape, and influence even the most well-studied genetic pathways.

## MATERIALS AND METHODS

**Bacterial strains and plasmids:** The ∼976 kb plasmid pMPPla107 was originally described in *Pseudomonas syringae* strain MAFF301305 (also known as pv. *lachrymans* 107) by Baltrus et al. 2011. A draft assembly sequence for pMPPla107 can be found at Genbank accession <u>CM000959.1</u>. All strains and plasmids used in the study are listed in Table 1. The focal *P. stutzeri* strain in this paper, 23a24, was isolated from soil (Sikorski et al., 2002), and was chosen for its high competence for natural transformation. This strain has been selected and phenotypically modified in a variety of ways to carry out the necessary experiments, with these modifications listed in Table 1.

**Table 1.**
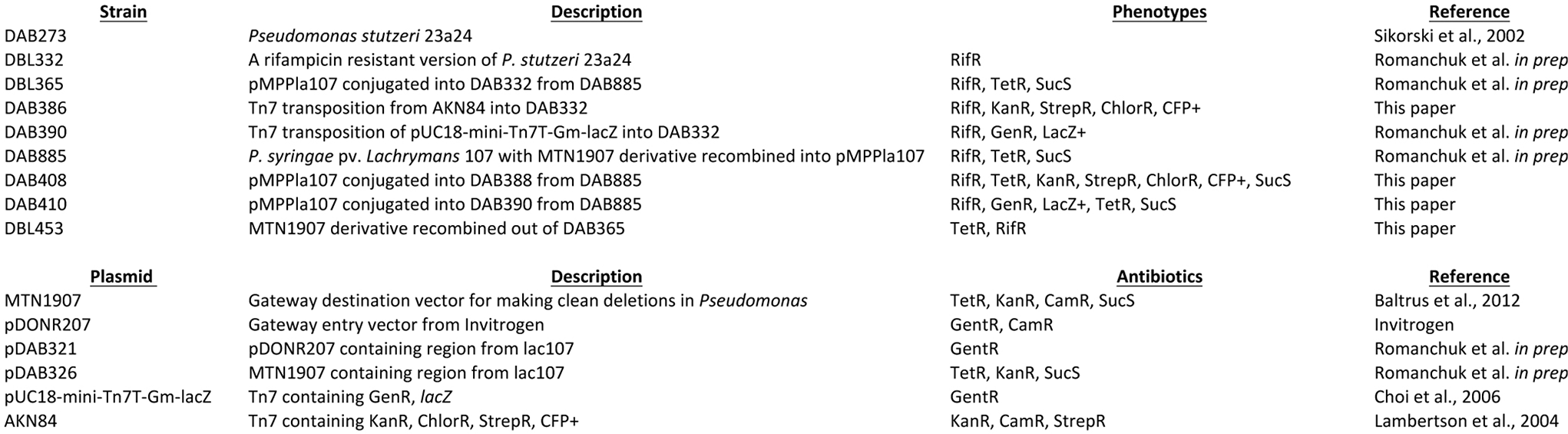
Strains and Plasmids

**Culture conditions:** All experiments were carried out at 27°C unless otherwise stated. Salt water LB (SWLB) was used as base media for liquid cultures and agar plates (Sikorski et al., 2002). All liquid cultures were incubated on a rotary shaker (200 rpm). Antibiotics were used at the following concentrations: 20 μg/mL tetracycline, 50 µg/mL kanamicin, 50 µg/mL rifampicin, 10µg/mL streptomycin. Xgal was used at a concentration of 40µg/mL. For competitive fitness experiments in the presence of naldixic acid, a concentration of 4µg/mL was used. All dilutions and cell suspensions took place in sterile 10 mM MgCl2.

**Competitive Fitness Assays:** To quantify the fitness cost of pMPPla107 in multiple environments, we set up independent competitive fitness assays, as described in Baltrus et al. 2008, for paired *P. stutzeri* strains (i.e., one that lacked (DBL332) & one that contained (DBL365) a megaplasmid tagged with tetracycline resistance) using a pMP-free control strain (DBL390) that was phenotypically marked with gentamycin resistance and lacZ. Each week, we revived the strains to LB agar plates and then SWLB liquid (containing tetracycline at 10µg/mL for pMP strains) by inoculating overnight cultures with cells from the corresponding agar plate. Subsequently, because the pMP impairs growth (e.g., extends the length of lag phase), we normalized the growth phase among the strains by conditioning each strain in tet-free SWLB for 48 hours under competition conditions by inoculating 2 mL cultures with 5 uL of the revival culture and incubating at 27°C. Furthermore, because the cost of pMPPla107 is severe at elevated temperatures and in the presence of nalidixic acid, we skewed the initial ratio of the pMP strain relative to DBL390 (∼5:1 test:control ratio), whereas the pMP-free strain was set up ∼1:1 with DBL390. Briefly, using optical density (OD600) to normalize cell density, we made a competition master mix (MM) for each strain large enough to make eight replicate 2 mL competitions per environment (e.g., 60 mL for 8 replicate competitions in three environments). However, because Nal 4 µg/mL was one of the environments tested, we initially set up the master mix with 2X the cells per unit volume (e.g., cells for 60 mL in 30 mL) and diluted one third of the master mix 50:50 with Nal 8 (i.e., 10 mL MM & 10 mL Nal 8 SWLB). The remaining 20 mL of the MM was diluted with 20 mL Nal 0 and used for the 8-replicate 27C/Nal0 & 8-replicate 35C/Nal0 competitions. For each replicate competition, relative fitness was calculated as the ratio of the growth rate (i.e., number of doublings) of the strain tested (e.g., DBL332, DBL365) relative to that of DBL390 during direct competition. The number of doublings for each strain in each competition was calculated by comparing the final density (i.e., [CFU]) and ratio of each independent competition to the average initial estimate calculated from eight replicate serial dilutions of each MM.

**Thermal Tolerance Assays:** Strains DBL332 and DBL365 were each grown up overnight in four replicate liquid cultures at 27°C in SWLB. A dilution series was then created from each tube and plated on two different SWLB agar plates. One plate was incubated at 27°C while the other was incubated at 37°C. This assay was carried out with two independently created megaplasmid strains (DBL365 and DBL412). After 4 days, both plates were photographed. Competitive fitness assays were performed as described above, with the following adjustments. The first set of replicate cultures was incubated for two days at 27°C, while the second set of replicates was incubated for two days at 35°C. 35°C was chosen as the temperature for competitions because DBL365 and DBL453 can undergo at least one doubling in these assays at this temperature. Each competitive fitness assay experiment contained 8 replicate cultures, and a total of 4 independent competitive fitness experiments were carried out for a total of 32 measurements for each strain. Within the ANOVA, strain and temperature were fixed variables while experiment was treated as a random variable.

**Antibiotic Resistance Assays:** Strains DBL332 and DBL365 were each grown up overnight in liquid cultures at 27°C in SWLB. Each culture was then used to inoculate a 96-well plate containing varying concentrations of either naldixic acid or 7-hydroxycoumarin. This 96-well plate was placed into a Biotek Synergy-H1 plate reader, and incubated at 27°C with shaking for 24 hours. Every hour we took measurements of OD_600_ from the assay plate in order to create a curve representing antibiotic inhibition. We chose 24 hours at the measurement represented in Figs. 1A and B because the starting OD_600_ values for each strain are equal. This assay was repeated with two independently created megaplasmid strains (DBL365 and DBL408), although only results from DBL365 are shown. Competitive fitness assays were performed as described above, with the following adjustments. A second set of replicates was inoculated into SWLB containing 4 µg/mL naldixic acid. After two days, all cultures were plated out to SWLB agar plates containing Xgal, and the ratio of test/control strains was counted. Each competitive fitness assay experiment contained 8 replicate cultures, and a total of 3 independent competitive fitness experiments were carried out for a total of 24 measurements for each strain. Within the ANOVA, strain and antibiotic were fixed variables while experiment was treated as a random variable.

**Biofilm Assay:** Strains DBL332 and DBL365 were each grown up overnight into 2 mL liquid cultures at 27°C in SWLB. After one day, a second culture was inoculated at a 1:100 dilution in 2 mL from the initial overnight culture and a sterile micropipette tip (Rainin, 1-20uL) was placed into the culture. This culture is incubated 4 days. At this point the pipette tip was extracted and thoroughly rinsed with 10 mM MgCl2 into a 2 mL eppendorf tube containing 4 glass beads to remove biofilm from the tip. The tube was then shaken in a FastPrep machine for 20 seconds at 4 m/s. The dislodged bacteria were enumerated by plating a dilution series on SWLB, incubating at 27°C, and counting colonies after 3 days. Each assay experiment contained 4 replicate cultures, and a total of 4 independent competitive fitness experiments were carried out for a total of 16 measurements for each strain. Within the ANOVA, strain and population location were fixed variables while experiment was treated as a random variable.

**Motility Assay**: Strains DBL332 and DBL365 were each grown up overnight into 2 mL liquid cultures at 27°C in SWLB. One uL of this culture is then inoculated into the center of a plate containing 1/2 strength SWLB and 0.25% agar. These plates were then incubated at 27°C for 2 days, at which point pictures of each plate are scanned and the size of the motility halo is analyzed using ImageJ. Each assay experiment contained 8 replicate cultures, and a total of 5 independent competitive fitness experiments were carried out for a total of 40 measurements for each strain. Within the ANOVA, strain and was a fixed variable while experiment was treated as a random variable.

**Bacteriocin Assay:** *P. aeruginosa* strains with killing activity were grown overnight in LB liquid culture at 37°C with shaking. 2mL were placed in a 2mL Eppendorf tube and centrifuged at 10,000 rpm for 5 minutes. Supernatant was carefully filter sterilized into a fresh tube without disturbing the pellet. Target strains were grown overnight in LB at 27°C with shaking. A 1:100 dilution in LB was made using the target strain overnight and grown for 4hr. 300µL were taken from these cultures and used to inoculate 3mL of melted agar (0.4%). Inoculated agar tubes were mixed well and utilized as an overlay on desired medium plates. Let overlay solidify with plate lid on for about 10-20min. Spot overlay with extracted killing supernatant and let dry. Incubate plates at 27°C for 1-2 days.

## Acknowledgements

Funding was provided by startup funds to David Baltrus from the University of Arizona as well as a Faculty Seed Grant from the University of Arizona Foundation

